# Sucrose phosphorylase from *Alteromonas mediterranea*: structural insight into the regioselective α-glucosylation of (+)-catechin

**DOI:** 10.1101/2023.04.11.536264

**Authors:** Marine Goux, Marie Demonceaux, Johann Hendrickx, Claude Solleux, Emilie Lormeau, Folmer Fredslund, David Tezé, Bernard Offmann, Corinne André-Miral

## Abstract

Sucrose phosphorylases, through transglycosylation reactions, are interesting enzymes that can transfer regioselectively glucose from sucrose, the donor substrate, onto acceptors like flavonoids to form glycoconjugates and hence modulate their solubility and bioactivity. Here, we report for the first time the structure of sucrose phosphorylase from the marine bacteria *Alteromonas mediterranea* (*Am*SP) and its enzymatic properties. Kinetics of sucrose hydrolysis and transglucosylation capacities on (+)-catechin were investigated. Wild-type enzyme (AmSP-WT) displayed high hydrolytic activity on sucrose and was devoid of transglucosylation activity on (+)-catechin. Two variants, *Am*SP-Q353F and *Am*SP-P140D catalysed the regiospecific transglucosylation of (+)-catechin: 89% of a novel compound (+)-catechin-4′-O-α-D-glucopyranoside (CAT-4’) for AmSP-P140D and 92% of (+)-catechin-3′-O-α-D-glucopyranoside (CAT-3’) for AmSP-Q353F. The compound CAT-4’ was fully characterized by NMR and mass spectrometry. An explanation for this difference in regiospecificity was provided at atomic level by molecular docking simulations: *Am*SP-P140D was found to preferentially bind (+)-catechin in a mode that favours glucosylation on its hydroxyl group in position 4’ while the binding mode in AmSP-Q353F favoured glucosylation on its hydroxyl group in position 3’.

## 1. Introduction

Oceans cover more than 70% of Earth’s surface and provides a unique environment to marine bacteria (*i.e.* high salinity, high pressure, low temperature and special lighting conditions). For decades, enzymes have been isolated and purified from terrestrial microorganisms, animals and plants. With the advent of biotechnology, there has been a growing interest and demand for enzymes with novel properties and robust biocatalysts. Due to its complexity, the marine environment represents a great opportunity for exploration of new enzymes and molecules [1, 2]. Marine enzymes are capable of being active under extreme conditions, which provide competitiveness and efficiency to different industrial processes [3, 4]. Among those, sucrose-phosphorylases (SPs) from the Glycoside Hydrolase family 13 subfamily 18 (GH13_18, EC 2.4.1.7) attract biotechnological interest as biocatalysts. Requiring only a cheap and abundant donor, SPs can perform transglucosylation reaction by transferring glucose from sucrose to an acceptor to yield α-glucosylated products with a retaining mechanism *via* a β-glucosyl-enzyme intermediate. Particularly, *Bifidobacterium adolescentis* SP (*Ba*SP-WT) and its mutants have been studied for biocatalytic synthesis of rare disaccharides [5, 6] and α-glucosylation of polyphenols [7, 8, 9, 10]. To date, the only documented structures of SP in the literature are from *Bifidobacterium adolescentis* [11]. Other SPs, from *Leuconostoc mesenteroides*, *Streptococcus mutans, Lactobacillus acidophilus* and *Thermoanaerobacterium thermosaccharolyticum*, have also been studied for the glucosylation of phenolic compounds [12, 13, 14]. Glucosylation of this type of molecules increases their solubility in water and their bioavailability in health, nutraceuticals and cosmetics applications [15, 16]. Controlling the regioselectivity of this glucosylation is also at stake for the synthesis of new compounds. We recently documented the activity of two variants of *Ba*SP-WT with respect to their ability to transfer regioselectively a glucose moiety onto (+)-catechin as an acceptor substrate [10]. *Ba*SP-Q345F and *Ba*SP-P134D/Q345F glucosylated (+)-catechin on hydroxyl groups in position 3’ (OH-3’) and 5 (OH-5) with obtainment of three glucosylated regioisomers: (+)-catechin-3′-O-α-D-glucopyranoside (CAT-3’), (+)-catechin-5-O-α-D-glucopyranoside (CAT-5) and (+)-catechin-3’,5-O-α-D-diglucopyranoside (CAT-3’,5), with a ratio of 51:25:24 for *Ba*SP-Q345F and 82:9:9 for *Ba*SP-P134D/Q345F.

*Alteromonas mediterranea*, also known as *Alteromonas macleodii* “Deep Ecotype” or AltDE, is an aerobic Gram-negative and mesophilic marine bacterium from the genus of *Proteobacteria* which was first isolated at a depth of 1000 m in the Eastern Mediterranean Sea in 2005 [17, 18, 19]. Wild type form of *Am*SP (*Am*SP-WT) shares 52% of global sequence identity with *Ba*SP-WT. Sequence alignments revealed that both enzymes possess highly conserved regions corresponding to the loop A (*Ba*SP-WT: ^336^AAASNLDLYQ^345^, *Am*SP-WT: ^344^AAASNLDLYQ^353^) and loop B (*Ba*SP-WT:^132^YRPRP^136^, *Am*SP-WT: ^138^FRPRP^142^) of the catalytic site [20]. In a preliminary screening using homology modelling and molecular docking, we identified that the catalytic cavity of the glucosyl-intermediate of *Am*SP-WT could potentially host a polyphenolic acceptor compound like (+)-catechin. Towards this end, the crystallographic structure of *Am*SP-WT was for the first time recently determined [Goux *et al.*, in preparation]. In the present work, we characterized *Am*SP-WT from a structural and functional perspective, and further analysed its structural and kinetic features. We also investigated the enzymatic properties of two variants of the enzyme towards their propensity to catalyse the regioselective transglucosylation of (+)-catechin. P140D and Q353F mutations, homologous to mutations P134D and Q345F of *Ba*SP-WT, displayed a single transfer reaction product for each enzyme: (+)-catechin-4′-O-α-D-glucopyranoside (CAT-4’) for *Am*SP-P140D and CAT-3’ for *Am*SP-Q353F. To explain the striking enzymatic activities of those variants, we provide in-depth structural insights by docking simulations and modelling. Our results interestingly broaden the available chemo-enzymatic synthetic tools for the efficient regioselective α-glucosylation of polyphenols.

## 2. Materials and methods

### 2.1. Chemicals

(+)-catechin was purchased from Extrasynthèse and PCR primers from Eurofins. All other chemicals were purchased from Sigma-Aldrich or VWR.

### 2.2. Vector construction and proteins

*Am*SP-WT and its variants were expressed as C-terminally hexahistidine-tagged proteins, allowing affinity purification by standard protocols. *Am*SP-WT gene (UniProt: S5AE64_9ALTE) was ordered from Genscript already cloned in a pET28b vector. Variants *Am*SP-P140D, *Am*SP-Q353F and *Am*SP-P140D/Q353F were obtained by site-directed mutagenesis using primers 1 and 2 for P140D mutation, and primers 3 and 4 for Q353F mutation (Table S1). *E. coli* BL21(DE3) competent cells (Novagen) were transformed and clones were selected using LB-agar medium supplemented with 25 μg/mL kanamycin and confirmed by Sanger sequencing (Eurofins Genomics). Proteins were produced, purified and characterized as previously described [10].

### 2.3. Kinetics of sucrose hydrolysis

All assays were performed in triplicate in 50 mM MOPS-NaOH at pH 8.0 in PCR strip tubes (Axygen) in a total volume of 100 µL at 25°C. Sucrose hydrolysis activities of *Am*SP WT (3 µM) and variants (10 µM) were obtained in the presence of sucrose as donor substrate ranging from 1 mM to 50 mM. At 0 and 40 min, 50 µL of reaction volume were sampled and inactivated at 95°C for 5 min. Glucose released due to sucrose hydrolysis was quantified by spectrophotometry thanks to the enzymatic coupled assay using glucose oxidase and peroxidase (GOD-POD), and 2,2’-azino-bis(3-ethylbenzothiazoline-6-sulfonic acid (ABTS) as chromogenic peroxidase substrate. In a 96-well plate, a volume of 25 µL of each sample were added to 200 µL of ABTS solution [5 mg/mL ABTS, 5 mg/mL glucose oxidase, 5 mg/mL peroxidase, 30 mM citrate buffer-NaOH pH 6.0]. Plates were incubated for 10 min at 25°C inside a Labsystems integrated EIA Management System and absorbance was measured at 405 nm. Kinetic data were fitted to Michaelis-Menten non-linear model and Hanes-Woolf linear model using respectively *nlm* and *lm* functions implemented in R to estimate the kinetic parameters (Figure S3 and S4, Table 1).

**Table 1:** Apparent kinetic parameters for sucrose hydrolysis by AmSP and its variants. Data were obtained by glucose titration using GOD/POD method of the reaction medium that reacted for 3h at 25°C and that contained the SP enzyme (10 µM for variants and 3 µM for WT) and 1 mM to 50 mM of sucrose, in MOPS-NaOH 50 mM pH 8.0 (n=3). Values are based on linear fit to the Hanes-Woolf model. Michaelis-Menten and Hanes-Woolf plots are provided in the ESI (Figures S3 and S4).

|  | 0% DMSO |  |  | 20% DMSO |  |  |
| --- | --- | --- | --- | --- | --- | --- |
| | $K_M$ (mM) | $k_{cat}$ (min<sup>-1</sup>) | $k_{cat}/K_M$ | $K_M$ (mM) | $k_{cat}$ (min<sup>-1</sup>) | $k_{cat}/K_M$ |
| *AmSP-WT* | 1.3 | 56.9 | 42.8 | 1.5 | 68.4 | 46.9 |
| *AmSP-P140D* | 2.2 | 14.7 | 6.7 | 1.0 | 24.3 | 24.1 |
| *AmSP-Q353F* | 7.1 | 3.0 | 0.4 | 2.3 | 5.6 | 2.5 |
| *AmSP-P140D/Q353F* | 12.6 | 2.1 | 0.2 | 6.9 | 1.1 | 0.2 |

### 2.4. Kinetics of the transglucosylation

Reactions were carried out in 50 mM MOPS-NaOH solution at pH 8.0 in a total volume of 1 mL. Reaction mixture containing 10 mM (+)-catechin in DMSO (1 eq., 100 μL), 20% DMSO (100 μL, (v/v)), 80 mM sucrose in H_2_O (8 eq., 100 μL) was incubated with a final concentration of 10 μM of purified enzyme at 25°C under slight agitation. Enzymatic synthesis was monitored by thin layer chromatography (TLC) for 24h. TLC plates were developed in solution composed of ethyl acetate/methanol/cyclohexane/water (7:1.5:1:1, v/v/v/v) with 0.1% formic acid (v/v). Products were visualized using a UV lamp at 254 nm and revealed with vanillin-sulphuric acid reagent. After centrifugation (ThermoScientific, Heraeus Pico17 Centrifuge, rotor 75003424, 20 min, 12 000g, 20°C), supernatant of the enzymatic reaction medium was analysed by analytical HPLC at 280 nm on a C-18 column (Interchim, 5µm, 250 x 4.6 mm, US5C18HQ-250/046) with an isocratic flow of 80% H2O, HCOOH 0.1% (v/v) and 20% MeOH, HCOOH 0.1% (v/v) for 20 min. Compound concentrations were calculated from the area under the curves obtained by analytical HPLC on a C-18 column. The assay was performed in triplicate.

### 2.5. Purification and analysis of glucosylated (+)-catechin

After 24h of incubation at 25°C under slight agitation, reaction media was centrifuged (12 000 *g*, 20 min) and supernatant was purified by HPLC at 280 nm on a C-18 column (Interchim, 5 μm, 250 x 21.2 mm, US5C18HQ-250/212) with a gradient system (solvent A: H2O HCOOH 0.1%; solvent B: MeOH, HCOOH 0.1%; t0 min = 70/30, t10 min = 70/30, t70 min = 10/90). Products were identified by mass spectroscopy (H^-^ mode, Waters, UPLC-MS2 high resolution) and NMR ^1^H and ^13^C in DMSO-d_6_ (Bruker, Pulse 400 MHz RS2D, 256 scans). Chemical shifts are quoted in parts per million (ppm) relative to the residual solvent peak. Coupling constant J are quoted in Hz. Multiplicities are indicated as d (doublet), t (triplet), m (multiplet). NMR peak assignments were confirmed using 2D ^1^H correlated spectroscopy (COSY), 2D ^1^H nuclear Overhauser effect spectroscopy (NOESY) and 2D ^1^H-^13^C heteronuclear single quantum coherence (HSQC). Complete characterization of CAT-4’ is provided in the ESI, for CAT-3’, CAT-5 and CAT-3’,5, please see [10].

### 2.6. Docking analysis of binding mode of (+)-catechin in the catalytic pocket

All docking experiments were performed with AutoDock Vina using, for each variant of the enzyme, 50 models of its glucosyl-intermediate form built using Rosetta suite of software [22] and

12 (+)-catechin conformers (see Electronic Supplementary Information for details of the modelling). Docking perimeter was limited to the residues of the active site of the enzyme. Each of the 12 conformers of (+)-catechin were docked on every conformer of the two variants of AmSP. This amounts to a total of 600 (50x12) docking experiments for each variant of the enzyme. Only the productive poses that could lead to a glucosylation of (+)-catechin were selected. To do so, docking poses were filtered using the following distance constraints: distances within 3.0 Å between any oxygen of (+)-catechin and the anomeric carbon atom C1 of the glucosyl moiety were assessed. Docking scores were compiled for these productive poses and compared between the two variants.

## 3. Results

### 3.1. Highly conserved structural features and potential activity of AmSP-WT

Through structural comparison of *Ba*SP-WT (PDB: 1R7A, 2GDV), *Ba*SP-Q345F (PDB: 5C8B), and our recently determined structure of *Am*SP-WT (PDB: 7ZNP), we identified the conserved residues likely involved in the reaction mechanism and potential substrate interactions (Table S2). The ‒1 subsite of SPs, also called donor site, has an optimal topology for binding glucose and is conserved between *Ba*SP-WT and *Am*SP-WT (Figure 1). The configuration of the two catalytic residues involved in the reaction mechanism are also almost identical (Figure 1A, in blue): a glutamyl residue acts as a general acid/base catalyst (E243 for *Am*SP-WT and E232 for *Ba*SP-WT) and an aspartyl residue performs the nucleophilic attack (D203 for *Am*SP-WT and D192 for *Ba*SP-WT). The third member of the catalytic triad (D301 for *Am*SP-WT and D290 for *Ba*SP-WT) stabilises the transition state with a strong hydrogen bond and presents also an identical configuration in the catalytic site. The structural elements that were shown to stabilize the glucosyl moiety in *Ba*SP-WT by non-polar contacts between a hydrophobic platform (F53/F156 for *Ba*SP-WT) and the hydrophobic C3-C4-C5 part of glucose are also conserved in *Am*SP-WT (F56/F167). The acceptor or +1 site of SPs is mainly shaped by two highly dynamic loops (Figure S1), which were shown to adopt different conformations based on the progress of the reaction mechanism: one conformation is the “donor binding mode” or closed conformation and the other is the “acceptor binding mode” or open conformation where an arginyl residue (R135 in *Ba*SP-WT) is thought to enable the enzyme to outcompete water as an acceptor through strong electrostatic interactions. When we compare the apoenzyme of *Ba*SP-WT (PDB: 1R7A) with our newly obtained apoenzyme of *Am*SP-WT (PDB: 7ZNP), a striking difference in the positioning of loop A is noticed. The enzyme has already an opened conformation and is in the “acceptor binding mode” with Y352 residue pointing inside the active site (Figure 1, in magenta) and R141 pointing outside (Figure 1, in orange). This open conformation was also observed for *Ba*SP-WT crystallized with the end-product of the reaction after hydrolysis of glucose (PDB: 2GDV, chain B). Moreover, crucial conserved residues involved in binding of both phosphate and fructose, Y196 and H234 for *Ba*SP-WT *vs.* Y207 and H245 for *Am*SP-WT, are in the same conformational positions thus allowing sucrose phosphorylase activity (Figure 1B, in cyan)[21]. In *Ba*SP-WT, Y132 is located at the entrance of the active site and contributes to sucrose specificity thanks to hydrophobic interactions with Y196 and F206. In *Am*SP-WT, the aromatic structure is conserved with the replacement of the tyrosinyl moiety by a phenylalanyl residue in position 138 (Figure 1B, in orange).

**Figure 1:**
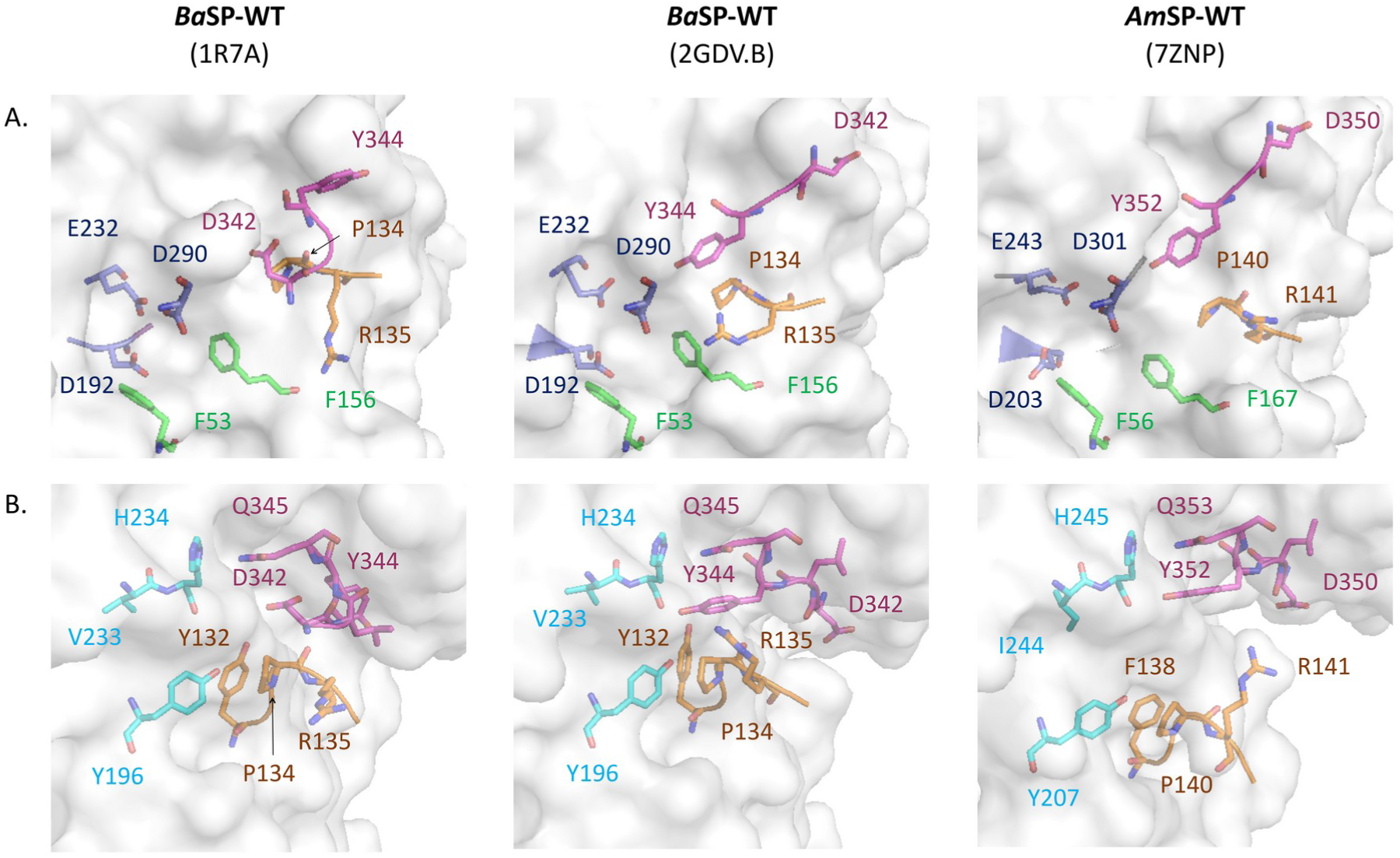
Crystallographic structures of *Ba*SP-WT and *Am*SP-WT focused on residues involved in (A) sucrose and (B) fructose binding. **(A)** In magenta: Loop A (in sticks for *Ba*SP-WT: Y344/D342, in sticks for *Am*SP-WT: Y352/D350); orange: Loop B (in sticks for *Ba*SP-WT: R135, in sticks for *Am*SP-WT: R141); green: hydrophobic platform (in sticks for *Ba*SP-WT: F53 and F156, in sticks for *Am*SP-WT: F56 and F167); and blue: residues of the catalytic triad (in sticks for *Ba*SP-WT: D192/E232/D290, in sticks for *Am*SP-WT: D203/E243/D301). **(B)** In magenta: Loop A (in sticks for *Ba*SP-WT: D342/L343/Y344/Q345, in sticks for *Am*SP-WT: D350/L351/Y352/Q353); orange: Loop B (in sticks for *Ba*SP-WT: Y132/R133/P134/R135, in sticks for *Am*SP-WT: F138/R139/P140/R141); Cyan: residues involved in sucrose phosphorylase activity (in sticks for *Ba*SP-WT: Y196/V233/H234, in sticks for *Am*SP-WT: Y207/I244/H245).

### 3.2. Determination of the apparent kinetic parameters for sucrose hydrolysis

C-terminally hexahistidine-tagged *Am*SP-WT and variants were produced in BL21(DE3) and purified by Ni-NTA immobilized metal affinity chromatography for further characterization. Interestingly, the P140D and Q353F mutations did not alter the enzyme stability, as evidenced by unchanged melting temperature (*T*_m_) of 43°C (Figure S2, Table S3). The apparent kinetic parameters for sucrose hydrolysis at 25°C were determined (Table 1). Globally, AmSP-WT has a higher apparent affinity towards sucrose (lower Km value) and higher turn-over number (kcat) than the studied variants. Lower activity for sucrose hydrolysis could be indicative of potential increased capacity in transglucosylation. For *Am*SP-WT, DMSO did not change significantly the *K*_m_ and *kcat* values. Interestingly, for the P140D and Q353F mono-variants, the presence of DMSO 20% increased the *kcat* two-fold and its specificity constant (*kcat*/*Km*) about 4-fold with respect to absence of co-solvent. With respect to the wild-type enzyme, in presence of DMSO, *Am*SP-P140D displayed a similar affinity for sucrose similar but a turnover number divided by almost 3-fold, leading to a kinetic efficiency (*kcat*/*Km*) divided by two. In the same conditions, *Am*SP-Q353F (Km = 2 mM) has a two fold decrease in affinity with respect to AmSP-WT (*K*_m_ =1 mM) and a corresponding 19-fold decrease in its turnover number.

### 3.3. (+)-catechin transglucosylation studies

We assessed the ability of SPs to transfer a glucose moiety from sucrose to (+)-catechin at 25°C after 24h under agitation (Figure S3). Observed products were purified by preparative HPLC and analysed by NMR and mass spectrometry. *Am*SP-P140D and *Am*SP-Q353F catalyse efficiently the synthesis of two different regioisomers of (+)-catechin glucoside: *Am*SP-P140D glucosylated mostly the hydroxyl groups in position 4’ (OH-4’) while *Am*SP-Q353F glucosylated the OH-3’ position (Figures S5B and S5C, Table S4). We monitored the synthesis of glucosylated (+)-catechin products during 24h by HPLC to determine conversion yields and proportion of regioisomers (Figure 2, Table 2). Interestingly, the main product formed with *Am*SP-P140D was CAT-4’ with a relative proportion of 89% while the main product formed with AmSP-Q353F was CAT-3’ with a relative proportion of 92%. The corresponding synthetic yields (percentage of (+)-catechin that was converted into these glycosylated products) was 26% and 82% respectively. These results clearly indicate that these two variants are highly regioselective with respect to their transglycosylation activities on (+)-catechin.

**Figure 2:**
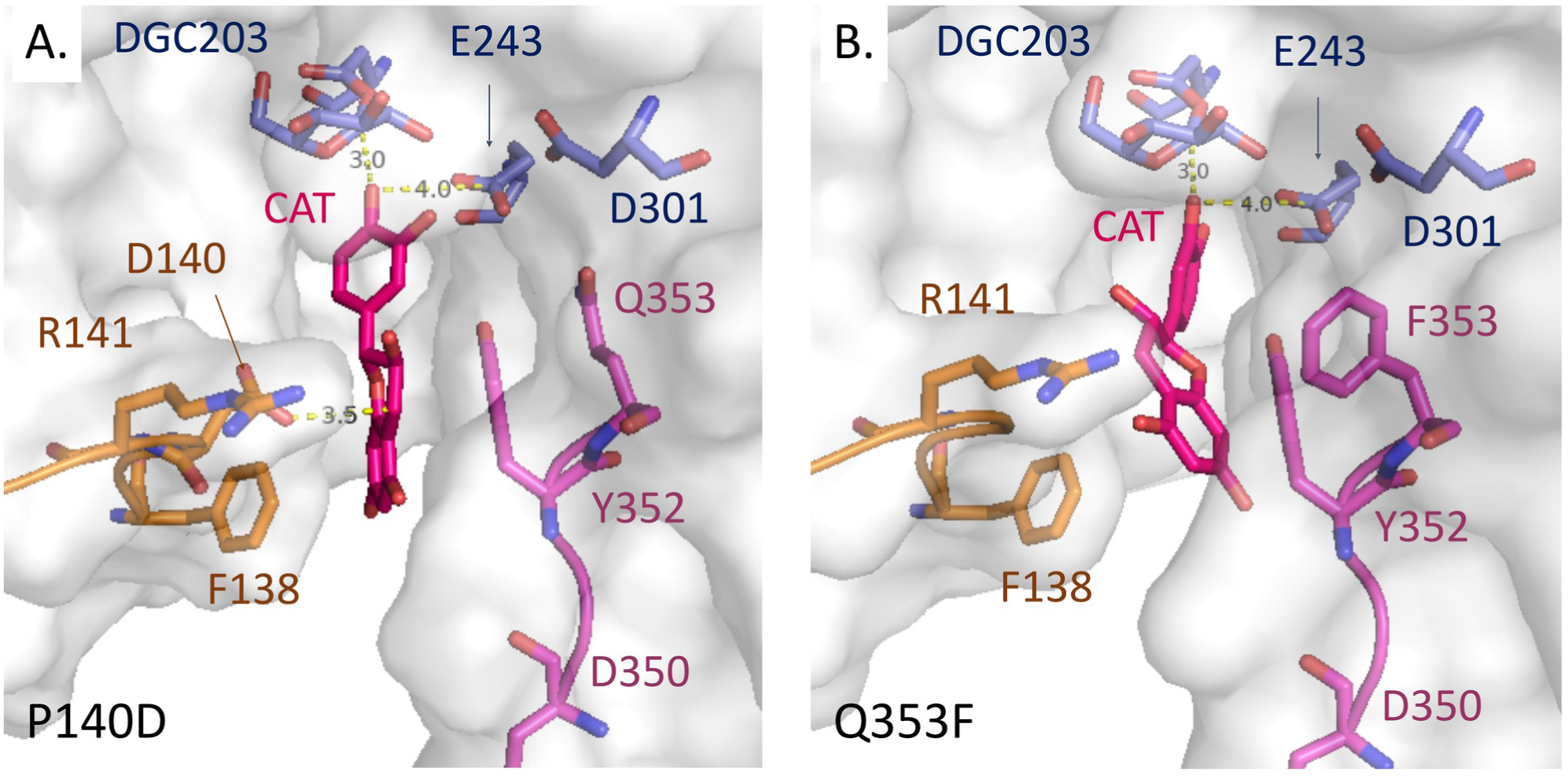
Products profile of AmSP-Q353F and AmSP-P140D using (+)-catechin as acceptor. Compound concentrations were calculated from the area under the curves obtained by analytical HPLC on a C-18 column for 7h and at 24 h of incubation at 25°C of the reaction mixture in a final volume of 1 mL in MOPS-NaOH 50 mM pH 8.0 with 10 µM of enzyme, 80 mM sucrose and 20% DMSO (v/v) (n=3). Errors correspond to standard deviations. Details of the data can be found in ESI (Table S4).

**Table 2:** Synthetic yields (A) and relative proportion (B) of (+)-catechin glycosylated products obtained with P140D and Q353F variants of AmSP. Synthetic yields and relative proportions of each product are expressed as a percentage of (+)-catechin that was converted into the corresponding glycosylated products (CAT-4’: (+)-catechin-4′-O-α-D-glucopyranoside; CAT-3’: (+)-catechin-3′-O-α-D-glucopyranoside; CAT-5: (+)-catechin-5-O-α-D-glucopyranoside; CAT3’,5: (+)-catechin-3’,5-O-α-D-diglucopyranoside). The relative proportion of each product was calculated from the area under the curves obtained by analytical HPLC at 24 h with the same conditions than for Figure S5. Values were obtained from the area under the curves obtained by analytical HPLC of the reaction mixture containing 10 μM enzyme, 10 mM (+)-catechin, 80 mM sucrose, 20% DMSO (v/v) in MOPS 50 mM pH 8.0, and incubated at 25°C for 24 h. Details of the data can be found in ESI (Table S4, Figure S5).

| Product | Synthetic yields |  |  |  | Proportion of each product |  |  |  |
| --- | --- | --- | --- | --- | --- | --- | --- | --- |
|  | CAT-4' | CAT-3' | CAT-5 | CAT-3',5 | CAT-4' | CAT-3' | CAT-5 | CAT-3',5 |
| **P140D** | 26% | 3% | Traces | Traces | 89% | 11% | Traces | Traces |
| **Q353F** | 3% | 83% | 2% | 2% | 4% | 92% | 2% | 2% |

### 3.4. Structural insights into the regioselectivity of AmSP-P140D and AmSP-Q353F

To further understand the observed regioselectivities, we performed molecular docking simulations on *Am*SP-Q353F and *Am*SP-P140D. The preferred orientations of (+)-catechin in the acceptor site of the glucosyl-enzyme intermediate were assessed. Docking results were consistent with the observed experimental regioselectivity. For *Am*SP-Q353F, binding mode of(+)-catechin suggest preference for transglycosylation in OH3’ position (Figures 3 and S7). On the other hand, for *Am*SP-P140D, the binding mode of (+)-catechin suggest preference for transglycosylation in OH-4’ position (Figures 3 and S7). For *Am*SP-P140D, (+)-catechin is stabilized in the active +1 site by a network of 4 hydrogen bonds and numerous hydrophobic contacts (Figure 4).

**Figure 3:**
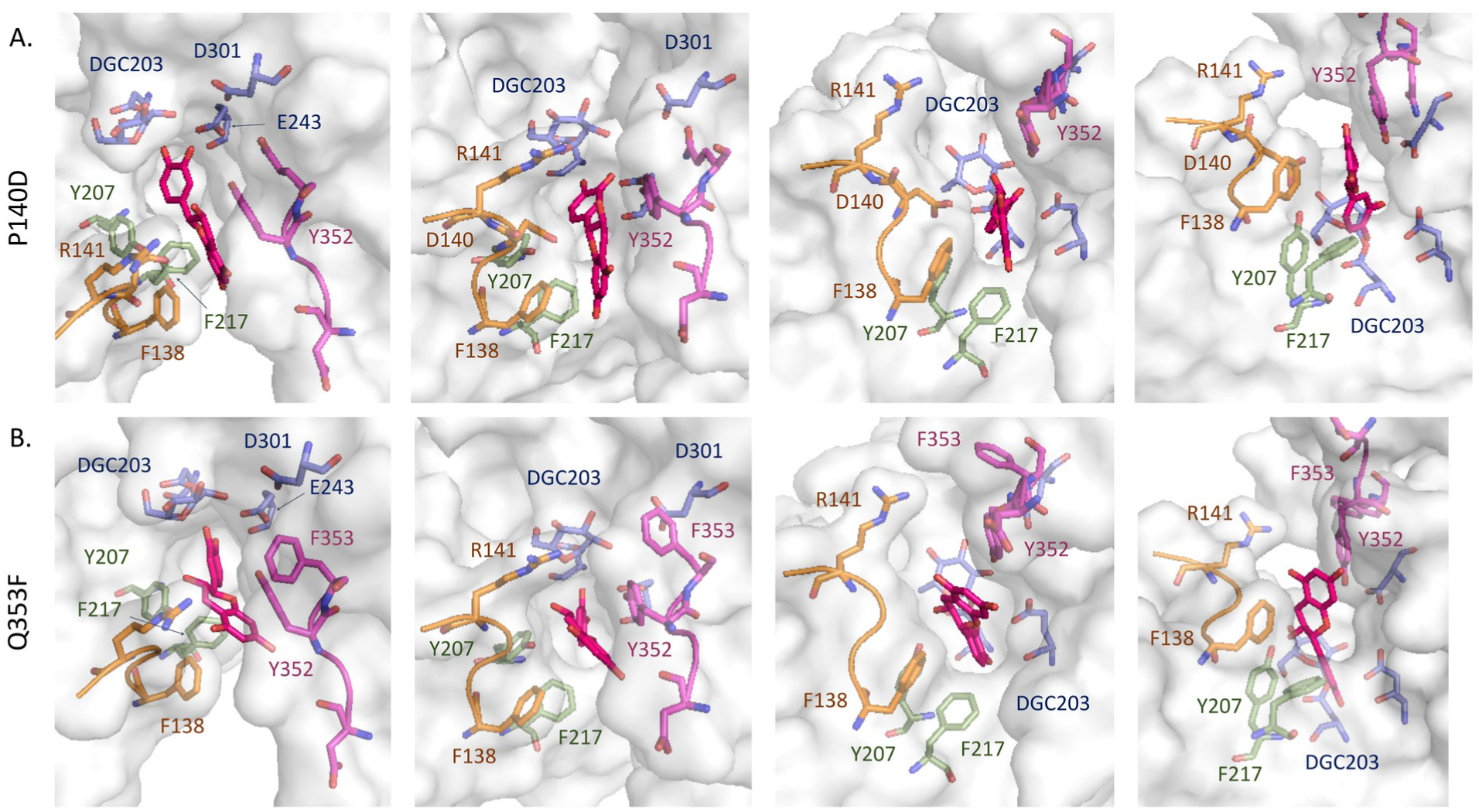
Comparison of the binding mode of (+)-catechin in AmSP-P140D and AmSP-Q353F. (A) Best binding mode of (+)-catechin in AmSP-P140D glucosyl-enzyme intermediate. (B) Best binding mode of (+)-catechin in AmSP-Q353F glucosyl-enzyme intermediate. In magenta, loop A with in sticks Y352/D350/Q(F)353 residues; in orange, loop B in orange with in sticks R141/F138 and D140 for AmSP-P140D; in blue, residues of the catalytic triad with in sticks DGC203/E243/D301; in pink, (+)-catechin.

**Figure 4.**
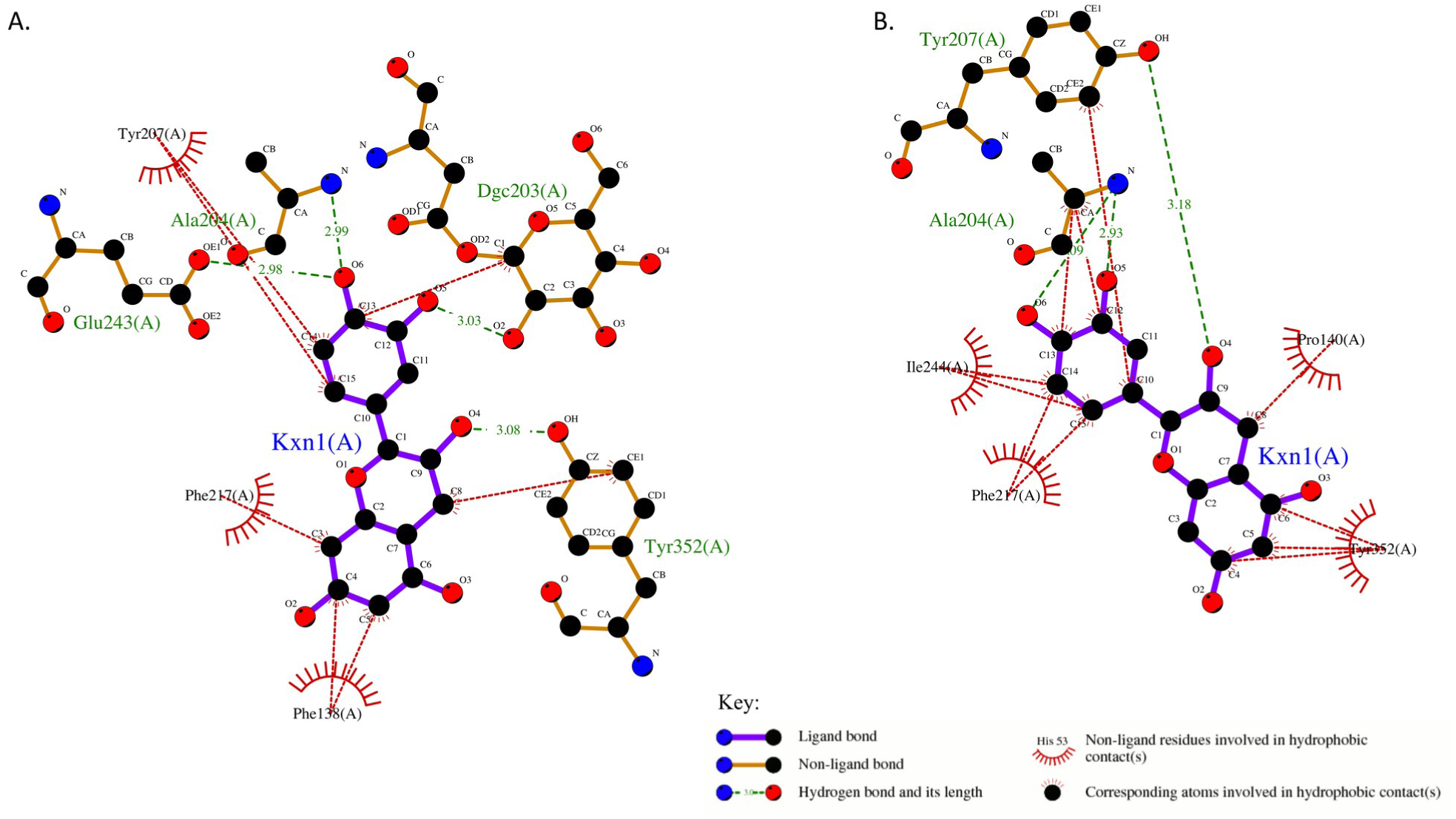
Analysis of the interaction between (+)-catechin and AmSP variants. Shown are hydrogen bonds and representative hydrophobic contacts between the (+)-catechin substrate (denoted Kxn1) and the interacting residues including the glucosylated-aspartyl (DGC203) in (A) AmSP-P140D and (B) AmSP-Q353F. The diagram was obtained using LigPlot Plus v.2.2 [24].

## 4. Discussion

In glycochemistry, the fine control of the regioselectivity is the Holy Grail in enzymatic reactions catalysed by glycosyl hydrolase (GHs) such as sucrose phosphorylase. A disadvantage of GHs is their moderate regioselectivity, meaning that a mixture of products is often formed when the acceptor contains more than one hydroxyl group. Previously, for (+)-catechin which consists of five phenolic hydroxyl groups, we generated with *Ba*SP-Q345F and *Ba*SP-P134D/Q345F a mixture of glucosylated regioisomers: CAT-3’, CAT-5 and CAT-3’,5 with a ratio of 51:25:24 for *Ba*SP-Q345F and 82:9:9 for *Ba*SP-P134D/Q345F. Another drawback is the relatively low product yields. With the same variants, we obtained a synthetic yield of 34%/15%/9% and 40%/5%/4%, respectively. In the active site of the SPs, the ‒1 site is rigid to allow a high selectivity on glucose while the +1 site is more flexible and can accept several types of acceptors or leaving groups. The +1 site of *Am*SP-WT, is mainly shaped by two highly labile loops, loop A (^344^AAASNLDLYQ^353^) and loop B (^138^FRPRP^142^), which undergo crucial conformational changes throughout the catalytic cycle suited for binding either fructose or phosphate. Crystal structure of *Am*SP-WT shows a wide access channel capable of accommodating naturally large polyphenolic acceptors. We decided to perform homologous mutations as what was done on *Ba*SP to check their impact on regioselectivity in the context of *Am*SP. By site-directed mutagenesis, we hence substituted the residue Q353 in loop A into F353 and/or the residue P140 into D140 in Loop B. We obtained three variants: *Am*SP-P140D, *Am*SP-Q353F and *Am*SP-P140D/Q353F. The double variant showed no improvement for the synthesis of CAT-3’ with the obtainment of a mixture of products at 25°C (Figure S5D) and was not further characterized.

By engineering only the residue 353 of the active site, we obtained an enhanced regioselectivity with glucosylation at OH-3’ position almost exclusively (Figure S5B). Interestingly, we also noticed that transglucosylation is most favored for this variant compared to hydrolysis in presence of (+)-catechin. Indeed, transglucosylation reaction proceeded to nearly completion within 24h (Figure 2) for AmSP-Q353F while rate of sucrose hydrolysis was very low (Table 1). *Am*SP-Q353F, as it was shown for *Ba*SP-Q345F and *Ba*SP-P134D/Q345F, preferentially glucosylate the OH-3’ position of flavonoids while ignoring the OH-4’ position. Docking studies confirmed that the most favoured pose for (+)-catechin in the catalytic +1 site of AmSP-Q353F lead to the formation of CAT-3’ (Figure 3A and 3C, and Table 2). As seen with BaSP-Q345F, we hypothesized that the introduction of F353 as a potential partner for π–π stacking would lead to rearrangements in loop A with a shift of Y352 which could stabilize (+)-catechin in the active site by hydrophobic interactions (Figures 4B and S8).

Surprisingly, while BaSP-P140D was not active, *Am*SP-P140D leads to the regioselective formation of CAT-4’ (Figure S5C and Table 2). Thus, the regioselectivity of *Am*SP shifted completely from CAT-3’ in Q353F variant to CAT-4’ in P140D variant. An explanation enlightened by molecular modelling is the steric hindrance caused by F138/Y207/F217 residues in AmSP-P140D that does not favor orientation of the substrate for CAT-3’ synthesis (Figures 3A, 3B and S8). Indeed, we observed that the conformation of the acceptor site drastically changed and seems to allow only an almost linear orientation of all three (+)-catechin rings (Figure S8). Contrarily to AmSP-Q353F, those very strong constraints lead to the regioselective glucosylation of the OH-4’ position of the flavonoid with a high proportion (89%). However, as shown in Figure 2, overall yield does not exceed 26% for the CAT-4’ transglucosylation product. We hypothesized that it is mainly due to the possible competition between (+)-catechin, sucrose and water into the active site, favoring the sucrose hydrolysis reaction (Figures S3 and S4).

## 5. Conclusion

In this study, we provided the first report of the use of variants of sucrose phosphorylase from *Alteromonas mediterranea* for the regioselective transglucosylation of (+)-catechin and the synthesis of a novel compound fully characterized, (+)-catechin-4′-O-α-D-glucopyranoside (CAT-4’). *Am*SP-Q353F and *Am*SP-P140D are able to synthesize regioselectively compound CAT-3’ and CAT-4’, with a proportion of 92% and 89%, respectively. With *Am*SP-P140D, we succeed to switch the regioselectivity from OH-3’ to OH-4’-glucosylated (+)-catechin. Mutation P140D changes drastically the conformation of the acceptor site and seems to allow an almost linear alignment of the glucose moiety and of all three (+)-catechin rings allowing selectively the glucosylation of the position OH-4’ of this flavonoid. Overall, the results described herein suggest that *Am*SP-Q353F and *Am*SP-P140D are suitable for the enzymatic regioselective synthesis of polyphenolic glucosides at high yields and could facilitate the synthesis of *de novo* products in OH-4’ position using other phenolic phytochemicals such as quercetin or kaempferol.

## Supporting information

Details about AmSP-WT sequence, primers, melting curves, kinetic parameters analysis, transglucosylation studies, HPLC/MS and NMR spectra are provided in the Electronic Supplementary Information.

## Author Contributions

MG and MD wrote the original draft. MG, MD, CM, BO and JH developed the methodology. MG, MD, CS and EL performed the experimental investigation and the subsequent analysis. DT and FF obtained the crystallographic structure of *Am*SP-WT. JH and BO performed the molecular and docking simulations and the following analysis. CM obtained the funding, designed and directed the project. All authors discussed the results and contributed to the final manuscript.

## Supporting information

Electronic Supplementary Information

## Acknowledgments

MG post-doctoral fellowship was supported by the “Region Pays de la Loire” and “Université Bretagne Loire” within the project “FunRégiOx”, and MD thesis by “Nantes Université”. We thank the CEISAM NMR platform for the NMR experiments.

