## Supplementary material for "Sucrose phosphorylase from *Alteromonas mediterranea*: structural insight into the regioselective α-glucosylation of (+)-catechin": Electronic Supplementary Information

#### Molecular modelling of variants of AmSP-WT

Glucosyl-enzyme intermediate 3D-models were built for AmSP-Q353F and AmSP-P140D using the following procedure and the Rosetta software [22]. Glucosylated-aspartyl 192 residue from chain A of crystal structure of BaSP-WT (PDB: 2GDV-A) was inserted into the crystal structure AmSP-WT (PDB: 7ZNP) that served as initial template for both variants. As this glucosylated aspartyl is a non-standard residue, it was absent from the database of the Rosetta software. Using Pymol3, the initial coordinates of this modified residue were retrieved. While this residue (D192) and the glucose moiety (BGC) are covalently linked in the crystal structure, the Pymol software considered them as two distinct residues. Thus, they were merged them into a single non-standard residue, which was called with a new ID, DGC. Associated charges and rotamers were calculated for this new residue using the Rosetta software. All those data were merged a single file and were added into the Rosetta database (Section 2.7).

With the DGC residue ready to be used, glucosyl-intermediates were built for the two variants of AmSP-WT. From the crystal structure (PDB: 7ZNP), using Rosetta, the native aspartyl residue in position 203 was mutated by the glucosylated-aspartyl DGC residue together with either the P140D or Q353F mutation. For each variant (AmSP-Q353F or AmSP-P140D), a sample of 50 conformers was generated thanks to the program Backrub from Rosetta suite, with 10 000 tries. In parallel, 12 conformers of (+)-catechin were also generated using the Mercury software (CCDC) [23] from the crystal structure OZIDOR of (+)-catechin.

### Characteristics of *Am*SP and its variants

##### *Sequence of AmSP-WT including His-tag on C-term*

```
  1 MGSIRNGVQL ITYADRLGDG NIESLTNLLD GPLKGLFKGV HILPFYYPYD GEDAGFDPID
 61 HTTVDERLGD WNNIKKLGES VDIMADLIVN HMSGQSEAFT DVLKKGRESE YWPLFLTKED
121 VFSGNDQAEI DEQIAKVFRP RPTPFFSDYE VGIETDSTET VPFWTTFTSN QIDIDVESEL
181 GKEYLSSILQ SFTESNVDLI RLDAAGYAIK RAGSNCFMLE ETFEFIEALS KRARTMGMQC
241 LVEIHSHYQT QIDIAARCDS VYDFALPPLV LHTLFTKDAS ALAHWLSISP RNCFTVLDTH
301 DGIGIVDVGA SGDKPGLISA DAINALVEQI HVNSNGESKK ATGAAANNVD LYQVNCTYYD
361 ALGKDDFAYL VARAIQFFSP GIPQVYYGGL LAAHNDMELL ANTNVGRDIN RPYLTTAMVE
421 DAIQKPVVKG LMQLITLRNE NKAFGGAFDV TYTDNTLVLS WSNDGDAASL TVDFAAMDAT
481 INTVSNGEES TLSIGALLAH HHHHH
```

**Table S1: List of primers used for quickchange mutagenesis**

| | |
|---|---|
| Primer 1 | CAAATTGCGAAAGTTTTTCGTGATCGTCCGACCCCGTTCTTTAGC |
| Primer 2 | GCTAAAGAACGGGGTCGGACGATCACGAAAAACTTTCGCAATTTG |
| Primer 3 | CGAACAACGTGGACCTGTACTTTGTTAACTGCACCTACTATGATG |
| Primer 4 | CATCATAGTAGGTGCAGTTAACAAAGTACAGGTCCACGTTGTTCG |

#### ***Structural homology between BaSP and AmSP***

A.

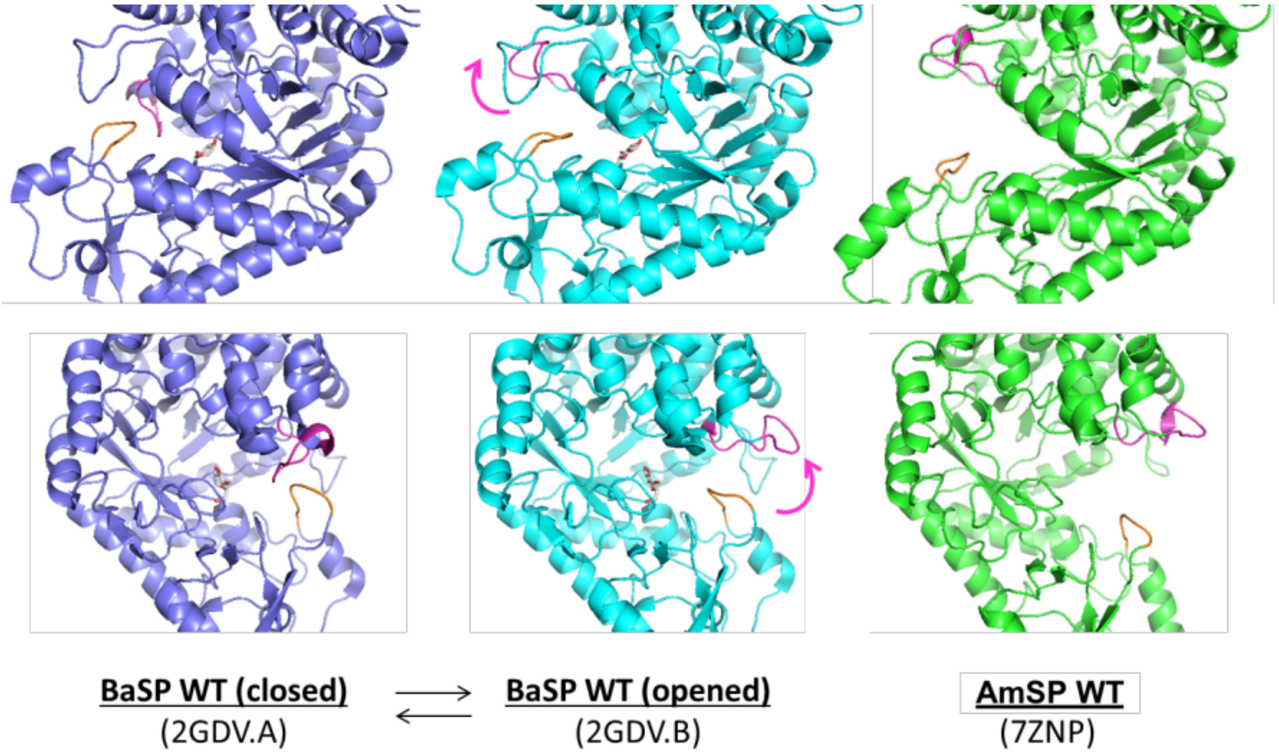

B.

| | | | | | | | | | | | |
|---|---|---|---|---|---|---|---|---|---|---|---|
| *Ba*SP-WT | 336 | A | A | A | S | N | L | D | L | Y | Q |
| *Am*SP-WT | 344 | A | A | A | S | N | L | D | L | Y | Q |

C.

| | | | | | | |
|---|---|---|---|---|---|---|
| *Ba*SP-WT | 132 | Y | R | P | R | P |
| *Am*SP-WT | 138 | F | R | P | R | P |

**Figure S1: Structural homology between *Ba*SP-WT and *Am*SP-WT. (A)** Comparison of closed (purple, PDB: 2GDV.A) and opened (cyan, PDB: 2GDV.B) conformations of *Ba*SP-WT and structure of *Am*SP-WT (green, PDB: 7ZNP). The functional loops are featured in magenta for loop A (involved in sucrose binding) and in orange for loop B (involved with polyphenol binding). **(B)** Sequence comparison for loop A between *Ba*SP-WT and *Am*SP-WT. **(C)** Sequence comparison for loop B between *Ba*SP-WT and *Am*SP-WT. ~~*: these residues are involved in the binding of phosphate.~~

**Table S2: Conserved residues and potential substrate interaction of *Am*SP.** Potential substrate interaction and function of conserved residues were determined using structural homology between crystal structures of apoenzyme *Am*SP-WT (PDB: 7ZNP), apoenzyme *Ba*SP-WT (1R7A) and open conformation BaSP-WT (2GDV.B). Residue numbering of *Am*SP derived from crystal structure 7ZNP. Residue numbering of BaSP came from crystal structures 1R7A and 2GDV.B. Numbered OH-groups and C-atoms address to sucrose (apostrophe for fructosyl moiety), unless stated otherwise. [a]: residues from the -1 subsite, [b]: residues from the Loop A/+1 subsite, [c]: residues from the Loop B/+1 subsite. Approx.: Approximately the same position.

| *Am*SP | *Ba*SP | Conserved Conformation 1R7A/2GDV.B | Potential substrate interaction/remarks | Potential function |
|---|---|---|---|---|
| **Asp203** [a] | Asp192 | Yes | Hydrogen bond with OH6 | catalytic nucleophile |
| **Glu243** [a] | Glu232 | Yes | Hydrogen bond with OH1 and OH1’ | general acid/base catalyst |
| **Asp301** [a] | Asp290 | Yes | Hydrogen bond with OH2 | transition state stabiliser |
| **Phe56** [a] | Phe53 | Yes | hydrophobic/π interaction with C3-C4-C5; Cation-π interaction with oxocarbenium ion-like transition state | hydrophobic platform |
| **Phe167** [a] | Phe156 | Yes | hydrophobic/π interaction with C6 and C1’ | hydrophobic platform |
| **His91** [a] | His88 | Approx. | Hydrogen bond with OH6 | binding of glycosyl moiety |
| **Arg201** [a] | Arg190 | Yes | Hydrogen bond with OH2 | binding of glycosyl moiety |
| **His300** [a] | His289 | Yes | Hydrogen bond with OH2 and OH3 | binding of glycosyl moiety |
| **Asp53** [a] | Asp50 | Yes | Hydrogen bond with OH4 | binding of glycosyl moiety |
| **Arg407** [a] | Arg399 | Yes | Hydrogen bond with OH3 and OH4 | binding of glycosyl moiety |
| **Gln171** | Gln160 | Approx. | Hydrogen bond with OH6 | binding of glycosyl moiety |
| **Ala204** | Ala193 | Yes | hydrophobic interaction with C6 and C1’ | binding of glycosyl/fructosyl moiety |
| **Leu351** [b] | Leu341 | No | hydrophobic interaction with C6 and C6’ | binding of glycosyl/fructosyl moiety |
| **Tyr207** | Tyr196 | Approx. | hydrophobic/π interaction with C1’ | involved in fructose binding |
| **Asp350** [b] | Asp342 | No/Approx. | Hydrogen bond with OH4’ | involved in fructose binding |
| **Gln353** [b] | Gln345 | Yes | Hydrogen bond with O3’ and O6' | involved in fructose binding |
| **Phe138** [c] | Tyr132 | Approx. | at the entrance of active site; hydrophobic surroundings | Specificity for fructose |
| **Pro140** [c] | Pro134 | Approx. | in vicinity of C4-OH of fructose | involved in the binding of the fructose-bound phosphate group |
| **Tyr207** | Tyr196 | Approx. | CE2-atom close to C1 of fructose | Specificity for fructose/phosphate |
| **His245** | His234 | Yes | in vicinity of C3 and C3-OH of fructose | Specificity for fructose/phosphate |
| **Arg141** [c] | Arg135 | No/Approx. | Hydrogen bond with phosphate O; Specificity for fructose/phosphate | involved in phosphate binding |
| **Leu351** [b] | Leu343 | No/Yes | in between important residues Asp342 and Tyr344 | Specificity for phosphate |
| **Tyr352** [b] | Tyr344 | No/Yes | Hydrogen bond with phosphate O | involved in phosphate binding |
| **Ile244** | Val233 | Yes | next to the acid/base catalyst; side chain turned away from active site | unknown |

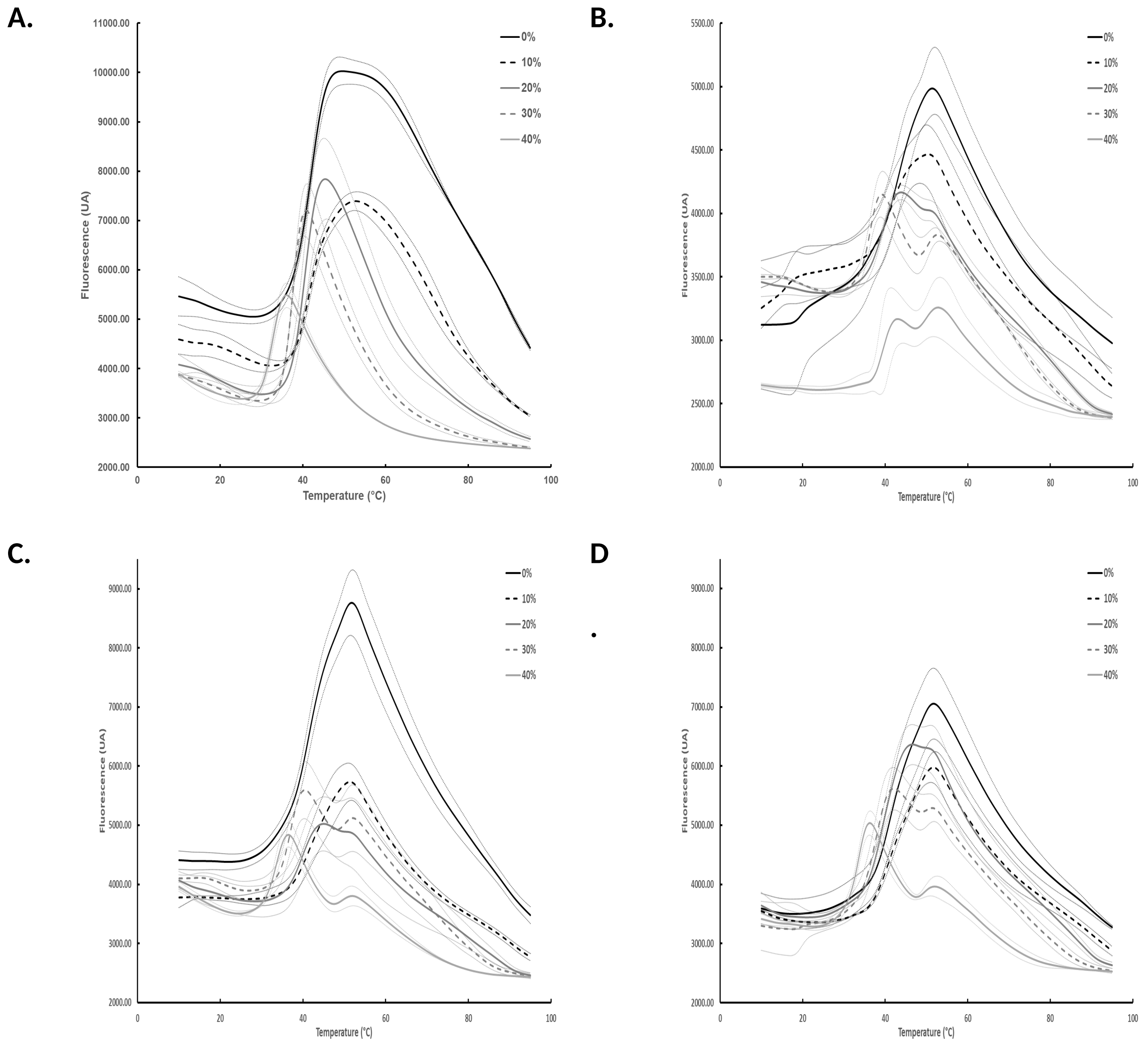

**Figure S2: Effect of DMSO on thermal stability of AmSP and its variants: (A) *Am*SP-WT, (B) *Am*SP-P140D, (C) *Am*SP-Q353F, (D) *Am*SP-P140D/Q353F.** Representative melting curve from 0% to 40% of DMSO (v/v) were obtained in a final volume of 25 µL in MOPS-NaOH 50 mM pH 8.0 with 3 µM of enzyme and Sypro orange 5x (n=3). The multiphase curve in presence of DMSO suggests the presence of a multi-domain protein or protein aggregation. Compared to the single curve obtained without DMSO, increasing DMSO concentration is shown to increase the destabilization of the enzymes.

**Table S3: Melting temperature of *Am*SP and its variants.** Values, given in °C, were obtained by calculating the first derivative -(dRFU)/dT of the melting curves.

| | WT | P140D | Q353F | P140D/Q353F |
|---|---|---|---|---|
| **$H_2O$** | 42.0 ± 0.0 | 42.8 ± 0.3 | 42.0 ± 0.9 | 42.2 ± 0.3 |
| **NPI-5** | 42.7 ± 0.3 | 42.5 ± 0.0 | 42.2 ± 0.3 | 41.5 ± 0.0 |
| **NPI-250** | 38.7 ± 0.3 | 38.2 ± 0.3 | 38.3 ± 0.3 | 37.3 ± 0.3 |
| **MOPS pH 7.0** | 44.8 ± 0.6 | 45.2 ± 0.3 | 44.0 ± 0.0 | 43.7 ± 0.3 |
| **pH 7.0 10%D** | 42.5 ± 0.0 | 42.2 ± 0.3 | 41.8 ± 0.3 | 42.3 ± 0.3 |
| **pH 7.0 20%D** | 39.5 ± 0.0 | 39.0 ± 0.0 | 39.2 ± 0.3 | 39.5 ± 0.0 |
| **pH 7.0 30%D** | 36.3 ± 1.0 | 36.3 ± 0.3 | 36.5 ± 0.5 | 36.7 ± 0.3 |
| **pH 7.0 40%D** | 31.3 ± 0.3 | 32.5 ± 0.0 | 32.7 ± 0.3 | 33.2 ± 0.3 |
| **MOPS pH 8.0** | 42.0 ± 0.0 | 43.0 ± 0.0 | 41.5 ± 0.0 | 41.2 ± 0.3 |
| **pH 8.0 10%D** | 41.3 ± 0.3 | 41.3 ± 0.4 | 41.5 ± 0.5 | 40.8 ± 1.0 |
| **pH 8.0 20%D** | 40.5 ± 0.0 | 39.3 ± 0.6 | 39.8 ± 0.3 | 39.5 ± 0.5 |
| **pH 8.0 30%D** | 37.5 ± 0.5 | 39.0 ± 0.7 | 37.0 ± 0.0 | 38.0 ± 0.5 |
| **pH 8.0 40%D** | 32.7 ± 0.3 | 39.5 ± 1.4 | 33.7 ± 0.3 | 33.7 ± 0.3 |
| **Citrate** | 45.5 ± 0.0 | 43.8 ± 0.4 | 43.5 ± 0.0 | 43.3 ± 0.3 |
| **HEPES** | 43.5 ± 0.3 | 44.0 ± 0.0 | 42.2 ± 0.6 | 42.2 ± 0.6 |
| **HEPES NaCl DTT** | 44.8 ± 0.6 | 45.0 ± 0.0 | 43.7 ± 0.8 | 44.3 ± 0.3 |

#### Determination of the apparent kinetic parameters

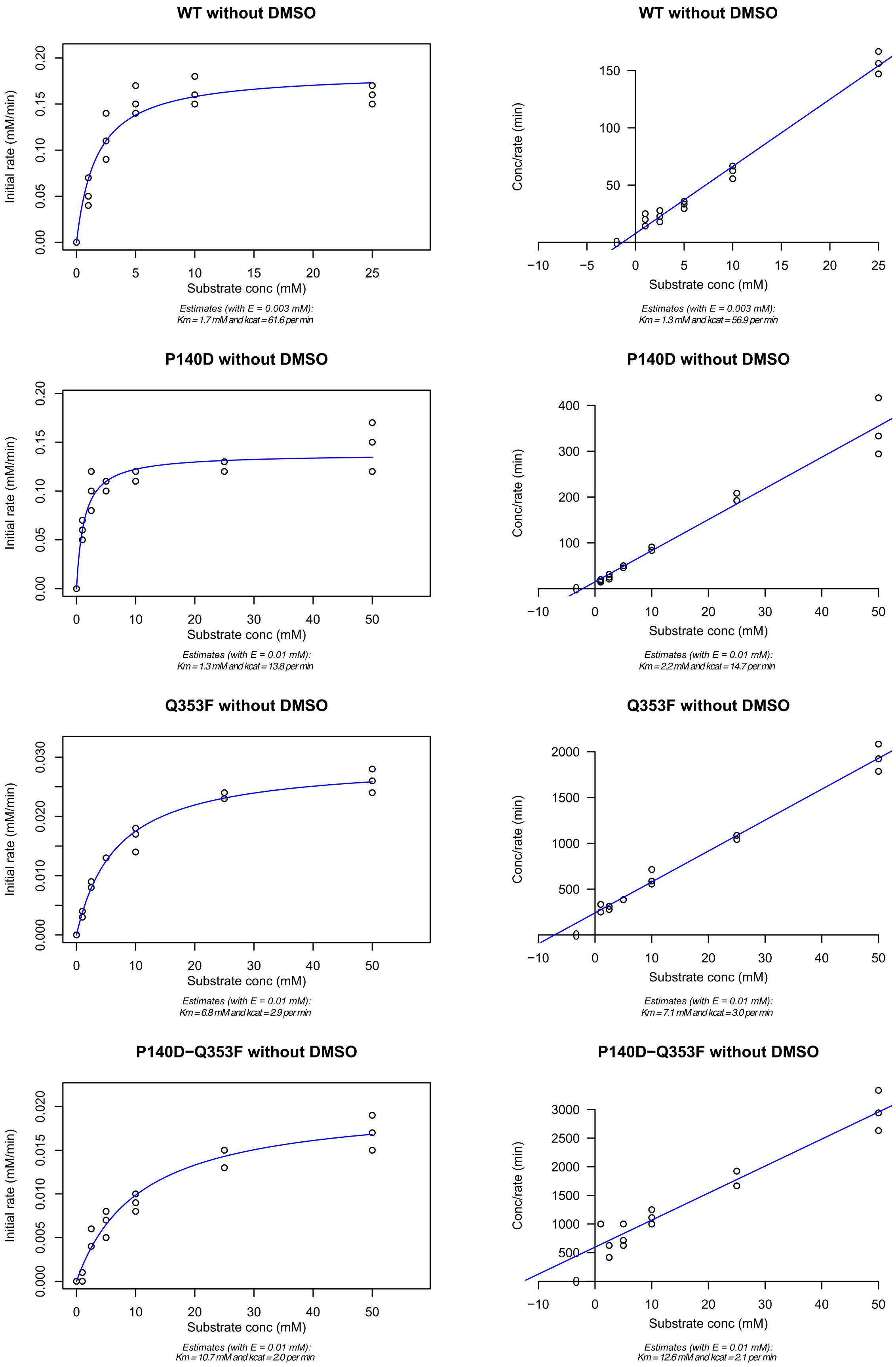

**Figure S3: Michaelis-Menten and Hanes-Woolf plots of sucrose hydrolysis in absence of DMSO obtained with *Am*SP-WT and its variants (P140D, Q353F and P140D/Q353F).** Data were obtained by glucose titration using GOD/POD method of the reaction medium that reacted for 3h at 25°C and that contained the SP enzyme (10 µM for variants and 3 µM for WT) and 1mM to 50 mM of sucrose, in MOPS-NaOH 50 mM pH 8.0 (n=3). Kinetic data were fitted to Michaelis-Menten non-linear model and Hanes-Woolf linear model using respectively *nlm* and *lm* functions implemented in R to estimate the kinetic parameters.

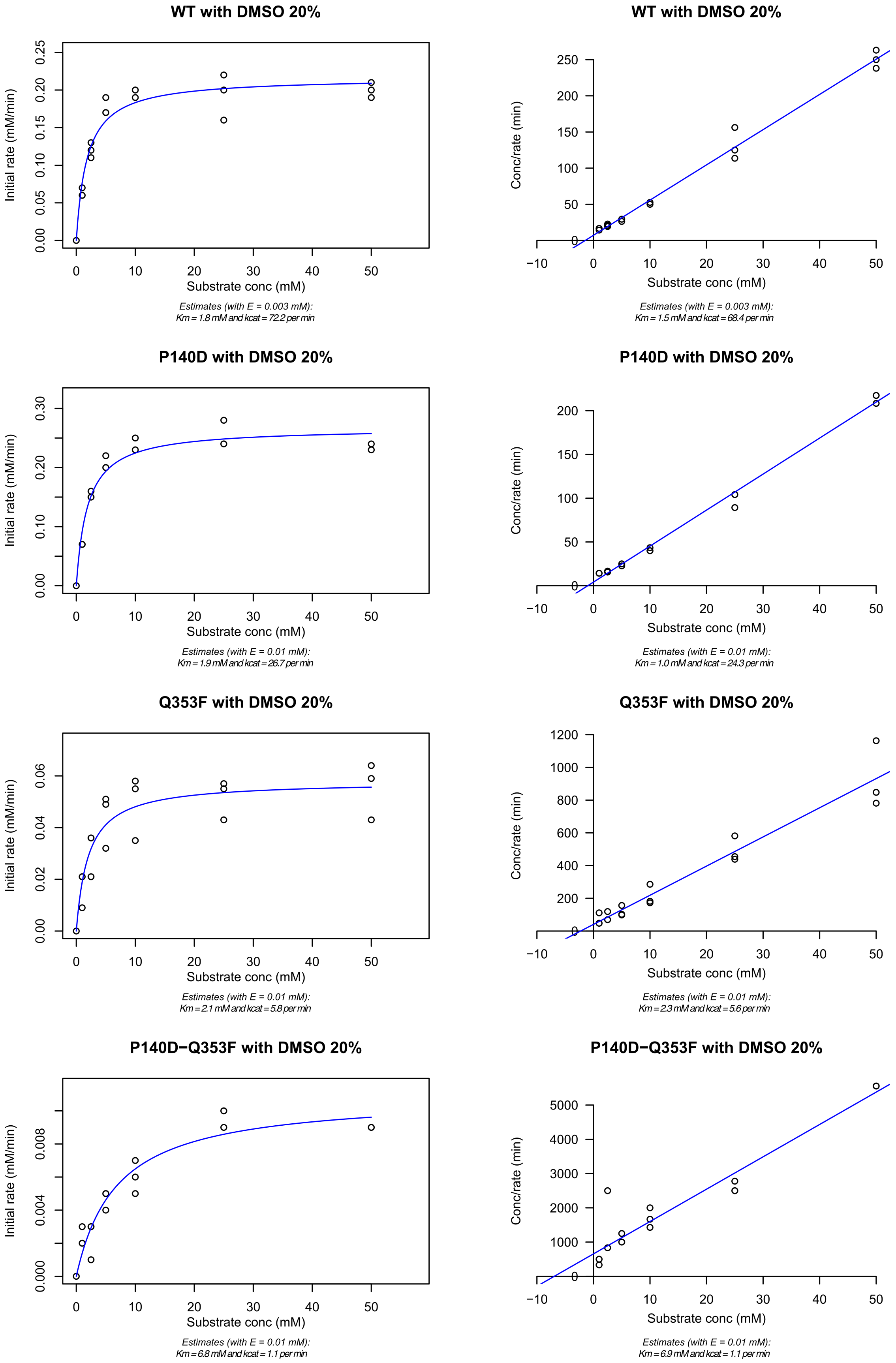

**Figure S4: Michaelis-Menten and Hanes-Woolf plots of sucrose hydrolysis obtained with *Am*SP-WT and its variants (P140D, Q353F and P140D/Q353F) in presence of 20% DMSO (v/v).** Data were obtained by glucose titration using GOD/POD method of the reaction medium that reacted for 3h at 25°C and that contained the SP enzyme (10 µM for variants and 3 µM for WT), 1mM to 50 mM of sucrose and DMSO 20% (v/v), in MOPS-NaOH 50 mM pH 8.0 (n=3). Kinetic data were fitted to Michaelis-Menten non-linear model and Hanes-Woolf linear model using respectively *nlm* and *lm* functions implemented in R to estimate the kinetic parameters

### (+)-catechin transglycosylation studies

#### *Synthesis yields for AmSP-Q353F and AmSP-P140D*

**Table S4: Compounds concentration obtained during (+)-catechin glucosylation by *Am*SP-Q353F and *Am*SP-P140D.** Compound concentrations were calculated from the area under the curves obtained by analytical HPLC on a C-18 column for 7 h and at 24 h of incubation at 25°C of the reaction mixture in a final volume of 1 mL in MOPS-NaOH 50 mM pH 8.0 with 10 µM of enzyme, 80 mM sucrose and 20% DMSO (v/v) (n=3).

***Am*SP-Q353F**

| Time (h) | CAT-3' (mM) | CAT-4' (mM) | CAT-5 (mM) | CAT-3',5 (mM) | (+)-catechin (mM) |
|---|---|---|---|---|---|
| 0 | 0.00 ± 0.00 | 0.00 ± 0.00 | 0.00 ± 0.00 | 0.00 ± 0.00 | 10.00 ± 0.00 |
| 1 | 1.12 ± 0.12 | 0.04 ± 0.00 | 0.04 ± 0.00 | 0.00 ± 0.00 | 8.80 ± 0.13 |
| 2 | 2.26 ± 0.22 | 0.08 ± 0.01 | 0.08 ± 0.01 | 0.00 ± 0.00 | 7.58 ± 0.23 |
| 3 | 3.23 ± 0.16 | 0.12 ± 0.01 | 0.11 ± 0.01 | 0.01 ± 0.00 | 6.53 ± 0.18 |
| 4 | 4.00 ± 0.14 | 0.15 ± 0.01 | 0.13 ± 0.01 | 0.01 ± 0.00 | 5.70 ± 0.16 |
| 5 | 4.74 ± 0.25 | 0.18 ± 0.02 | 0.16 ± 0.01 | 0.02 ± 0.00 | 4.90 ± 0.28 |
| 6 | 5.37 ± 0.21 | 0.20 ± 0.01 | 0.18 ± 0.01 | 0.03 ± 0.00 | 4.22 ± 0.23 |
| 24 | 8.36 ± 0.13 | 0.36 ± 0.01 | 0.21 ± 0.02 | 0.20 ± 0.02 | 0.88 ± 0.15 |

***Am*SP-P140D**

| Time (h) | CAT-3' (mM) | CAT-4' (mM) | (+)-catechin (mM) |
|---|---|---|---|
| 0 | 0.00 ± 0.00 | 0.00 ± 0.00 | 10.00 ± 0.00 |
| 1 | 0.05 ± 0.00 | 0.40 ± 0.02 | 9.55 ± 0.02 |
| 2 | 0.10 ± 0.00 | 0.74 ± 0.02 | 9.17 ± 0.02 |
| 3 | 0.12 ± 0.00 | 0.99 ± 0.02 | 8.88 ± 0.02 |
| 4 | 0.15 ± 0.00 | 1.19 ± 0.03 | 8.65 ± 0.03 |
| 5 | 0.17 ± 0.00 | 1.36 ± 0.06 | 8.47 ± 0.06 |
| 6 | 0.19 ± 0.01 | 1.49 ± 0.07 | 8.29 ± 0.03 |
| 24 | 0.31 ± 0.02 | 2.50 ± 0.19 | 7.19 ± 0.21 |

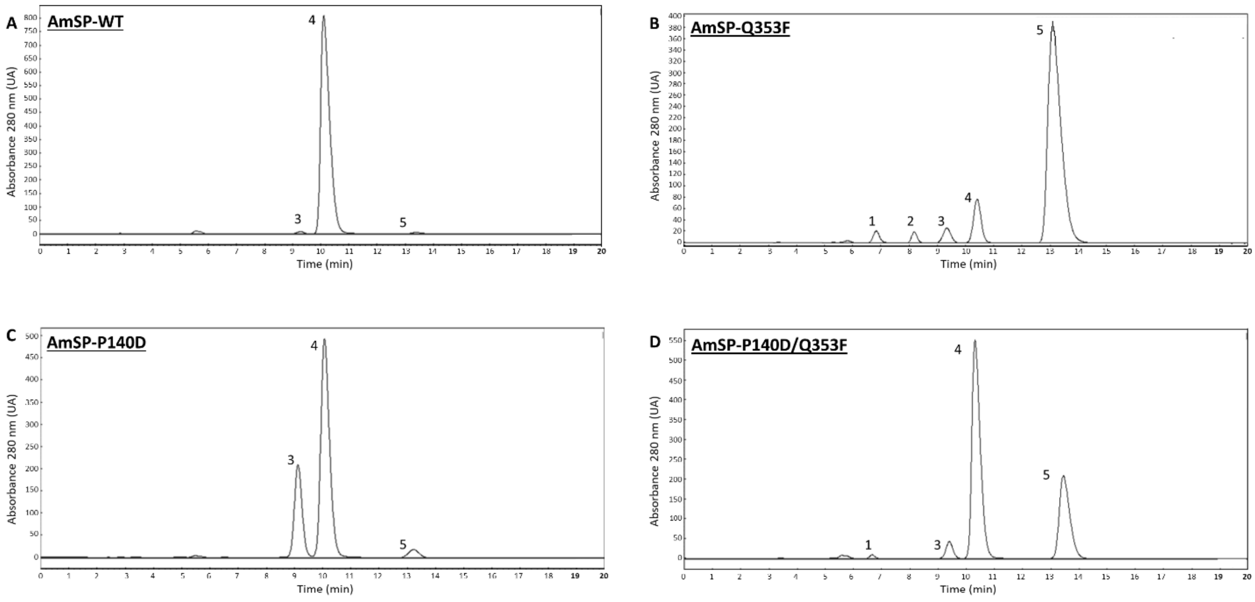

**Figure S5: HPLC chromatogram of a 24 h reaction medium of (+)-catechin glucosylation by *Am*SP-WT and its variants.** Peak 1: CAT-5, peak 2: CAT-3',5, peak 3: CAT-4', peak 4: (+)-catechin, peak 5: CAT-3'. Unassigned peaks correspond to impurities and degradation of (+)-catechin. (10 µM enzyme, 10 mM (+)-catechin, 80 mM sucrose, 20% DMSO (v/v) in MOPS 50 mM pH 8.0 at 25°C for 24 h). HPLC conditions: isocratic mode at 80% $H_2O$ (v/v), 0.1% formic acid (v/v) and 20% MeOH (v/v), 0.1% formic acid (v/v).

##### *$^{1}$H and $^{13}$C NMR Spectral Data of CAT-4' in DMSO*

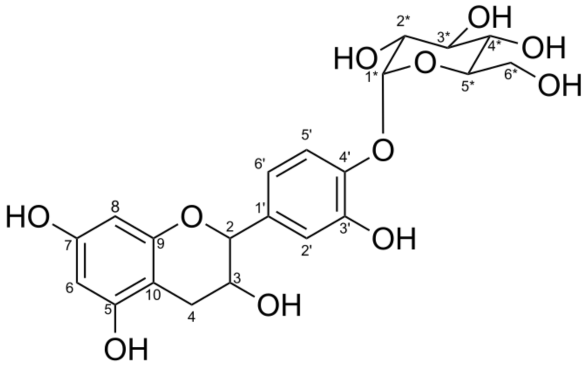

**MS (ESI positive):**

Ion Formula: $C_{21}H_{23}O_{11}$

m/z calculated: 451.1239

m/z experimental: 451.1240

error [ppm]: -0.2

$^{1}$H NMR (DMSO-$d_6$, δ) : 7.14 (d, $^{3}J_{5'-6'}$ = 8.3 Hz, 1H, $H_{5'}$), 6.80 (d, $^{3}J_{2'-6'}$ = 2.0 Hz, 1H, $H_{2'}$), 6.72 (dd, $^{3}J_{6'-2'}$ = 2.0 Hz, $^{3}J_{6'-5'}$ = 8.3 Hz, 1H, $H_{6'}$), 5.91 (d, $^{3}J_{6-8}$ = 2.2 Hz, 1H, $H_{6}$), 5.71 (d, $^{3}J_{8-6}$ = 2.2 Hz, 1H, $H_{8}$), 5.17 (d, $^{3}J_{1*-2*}$ = 3.6 Hz, 1H, $H_{1*}$), 4.55 (d, $^{3}J_{2-3}$ = 7.4 Hz, 1H, $H_{2}$), 3.85 (td, $^{3}J_{3-2}$ = 7.4 Hz, $^{3}J_{3-4a}$ = 5.3 Hz, $^{3}J_{3-4b}$ = 8.4 Hz, 1H, $H_{3}$), 3.71-3.65 (m, $^{3}J_{3*-2*}$ = 9.7 Hz, , 1H, $H_{3*}$), 3.65-3.62 (m, 1H, $H_{6a*}$), 3.55-3.49 (m, 1H, $H_{5*}$), 3.55-3.49 (m, 1H, $H_{6b*}$), 3.33 (dd, $^{3}J_{2*-1*}$ = 3.6 Hz, $^{3}J_{2*-3*}$ = 9.7 Hz, 1H, $H_{2*}$), 3.21-3.17 (m, 1H, $H_{4*}$), 2.65 (dd, $^{3}J_{4a-3}$ = 5.3 Hz, $^{2}J_{4a-4b}$ = 16.0 Hz, 1H, $H_{4a}$), 2.36 (dd, $^{3}J_{4b-3}$ = 8.4 Hz, $^{2}J_{4b-4a}$ = 16.3 Hz, 1H, $H_{4b}$).

$^{13}$C NMR (DMSO-$d_6$, δ) : 156.5 ($C_{7}$), 156.2 ($C_{5}$), 155.2 ($C_{10}$), 147.1 ($C_{4'}$), 144.7 ($C_{3'}$), 134.7 ($C_{1'}$), 118.2 ($C_{6'}$), 117.3 ($C_{5'}$), 114.8 ($C_{2'}$), 100.4 ($C_{1*}$), 99.0 ($C_{9}$), 95.2 ($C_{6}$), 93.8 ($C_{8}$), 80.7 ($C_{2}$), 73.8 ($C_{5*}$), 73.1 ($C_{3*}$), 72.0 ($C_{2*}$), 70.0 ($C_{4*}$), 66.3 ($C_{3}$), 60.7 ($C_{6*}$), 27.8 ($C_{4}$).

**Table S5: Comparison of $^{1}$H and $^{13}$C NMR Spectrum data of CAT-4’ and (+)-catechin in DMSO.** Multiplicity abbreviations for $^{1}$H NMR: d = doublet, dd = doublet of doublets, td = pseudo triplet of doublets, m = multiplet. For characterization of CAT-3’, CAT-5 and CAT-3’,5 please see [10].

| **POSITION** | **CATECHIN ∂1H; J (HZ)** | **CAT-4’ ∂1H; J (HZ)** | **CAT-4’ ∂13C** |
|---|---|---|---|
| **1** | X | X | X |
| **2** | 4.48 (d 7.4) | 4.55 (d 7.4) | 80.67 |
| **3** | 3.82 (td 8.1, 7.4, 5.2) | 3.85 (td 8.4, 7.4, 5.33) | 66.26 |
| **4** | 2.67 (dd 16.1, 5.2)<br>2.35 (dd 16.0, 8.1) | 2.65 (dd 16.0, 5.3)<br>2.36 (dd 16.3, 8.4) | 27.84 |
| **5** | X | X | 156.21 |
| **6** | 5.89 (d 2.2) | 5.91 (d 2.2) | 95.22 |
| **7** | X | X | 156.51 |
| **8** | 5.69 (d 2.2) | 5.71 (d 2.2) | 93.78 |
| **9** | X | X | 98.95 |
| **10** | X | X | 155.17 |
| **1'** | X | X | 134.68 |
| **2'** | *6.73 (d 1.9)* | *6.80 (d 2.0)* | 114.79 |
| **3'** | X | X | 144.71 |
| **4'** | X | X | 147.06 |
| **5'** | *6.69 (d 8.1)* | *7.14 (d 8.3)* | 117.26 |
| **6'** | *6.60 (dd 8.1, 1.9)* | *6.72 (dd 8.3, 2.0)* | 118.16 |
| **1*** | X | 5.17 (d 3.6) | 100.40 |
| **2*** | X | 3.33 (dd 9.7, 3.6) | 72.02 |
| **3*** | X | 3.71-3.65 (m) | 73.06 |
| **4*** | X | 3.21-3.17 (m) | 69.96 |
| **5*** | X | 3.59-3.55 (m) | 73.75 |
| **6*** | X | 3.65-3.62 (m)<br>3.55-3.49 (m) | 60.73 |

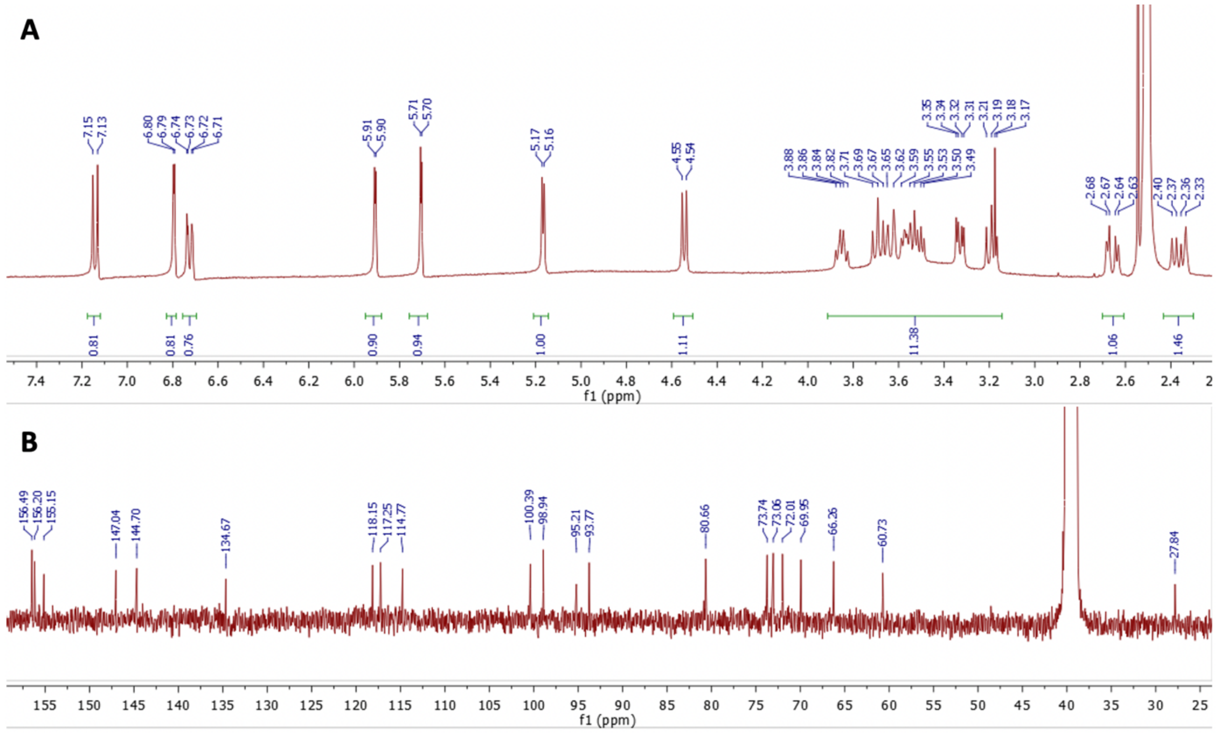

**Figure S6: NMR spectrum of (+)-catechin-4’-O-α-D-glucoside. (A)** $^{1}H$ NMR and **(B)** $^{13}C$ NMR in DMSO-d6 (400 MHz).

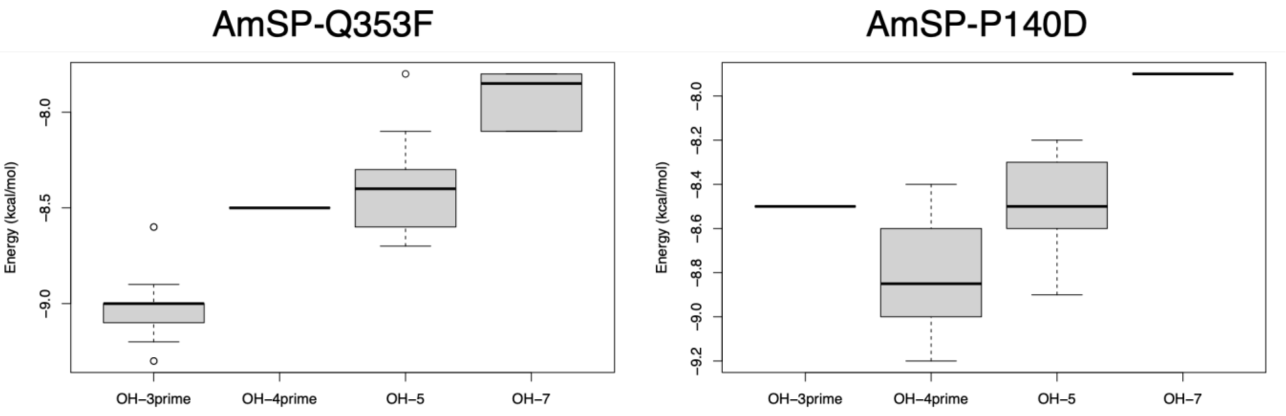

**Figure S7: Statistical analysis of the binding energies of productive poses for glucosylation of (+) catechin in OH-3', OH-4', OH-5 and OH-7 positions with Q353F and P140D.** Molecular docking poses were filtered by considering those with an oxygen of (+)-catechin within 3 Å of the C1 atom of the glucosyl moiety as productive (see sections 2.4, 2.5 and 3.4 in main text for additional details).

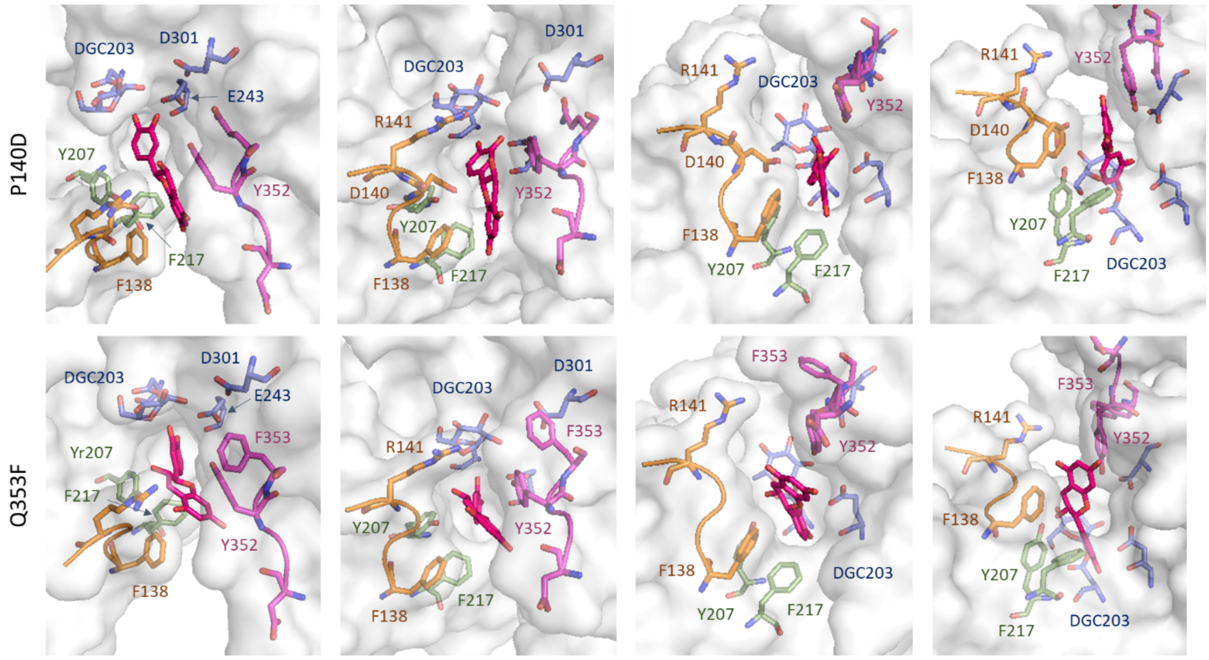

**Figure S8: Different views of the position of the best productive poses of (+)-catechin in the active site of AmSP-P140D and AmSP-Q353F.** In magenta: Loop A with in sticks Y352/F350/Q(F)353 residues; orange: Loop B with in sticks R141/F138 and D140 for AmSP-P140D; blue: residues of the catalytic triad with in sticks DGC203/E243/D301; green: aromatic residues involved in steric clash with in sticks Y207 and F217; and pink: (+)-catechin.
